# Modeling Metallicity: Low Level Visual Features Support Robust Material Perception

**DOI:** 10.1101/2021.02.22.432364

**Authors:** Joshua S. Harvey, Hannah E. Smithson

## Abstract

The human visual system is able to rapidly and accurately infer the material properties of objects and surfaces in the world. Yet an inverse optics approach—estimating the bi-directional reflectance distribution function of a surface, given its geometry and environment, and relating this to the optical properties of materials—is both intractable and computationally unaffordable. Rather, previous studies have found that the visual system may exploit low-level spatio-chromatic statistics as heuristics for material judgment. Here, we present results from psychophysics and modelling that supports the use of image statistics heuristics in the judgement of metallicity—the quality of appearance that suggests an object is made from metal. Using computer graphics, we generated stimuli that varied along two physical dimensions: the smoothness of a metal object, and the evenness of its transparent coating. This allowed for the manipulation of low-level image statistics, whilst ensuring that each stimulus was a naturalistic, physically plausible image. A conjoint-measurement task decoupled the contributions of these dimensions to the perception of metallicity. Low-level image features, as represented in the activations of oriented linear filters at different spatial scales, were found to correlate with the dimensions of the stimulus space, and decision-making models using these activations replicated observer performance in judging metal smoothness, coating bumpiness, and metallicity. Importantly, the performance of these models did not deteriorate when objects were rotated within their simulated scene, with corresponding changes in image properties. We therefore conclude that low-level image features may provide reliable cues for the robust perception of metallicity.

## Introduction

The twenty-first-century visual landscape is both littered and adorned with the unmistakable flash of metallic objects. Across architecture, fashion, food, transport, and technology, we are presented with more metallic surfaces than at any other time in history. Unlike other visual features, such as the reddening of ripening fruit, or the gloss of wettened surfaces, the human visual system did not evolve in an ecosystem of metallic stimuli. Nonetheless, we are highly accurate in our judgments of metallicity from vision alone.

Material perception may at first glance appear a simple and closed deductive problem; the material composition of objects will determine their surface reflectance properties, and hence the images received on the retina as a function of the object’s geometry and environmental illumination. But using images to estimate how light scatters off a surface as a function of all incoming and outgoing angles—the the bi-directional reflectance distribution function in graphics terminology—is intractable. Moreover, the visual system exhibits robustness across undetermined parameters, both of surfaces themselves and their surrounding environment. Metallicity, the visual quality of an object suggesting it is made of metal, is a prime example of this. The surfaces of most metallic objects return a solely specular (mirror) reflection to observers; their appearance is therefore highly contingent on both neighbouring emitters (light sources) and reflectors. At the same time, the physical smoothness and oxide-layer depths of metallic surfaces can result in appearances ranging anywhere from an immaculately polished mirror to the matte sheen of an anodized laptop. Throughout this range, and across viewing environments, we accurately judge such objects as metallic, rather than other shiny alternatives such as porcelain or glass^1–4^. At the same time, the recognition of metal can prove particularly problematic for computer vision systems^5,6^.

A sizable economic value is placed on the visual appearance of metallic surfaces in sectors such as the automotive and jewelry industries, and the literature on metallic appearance reflects this. There are numerous studies quantifying and perceptually evaluating the visual qualities of metallic surfaces and coatings, considering qualities such as colour, gloss^7^, scatter, brilliance, and lustre^8^, as well as ‘visual texture’ such as glitter, glint, and sparkle^9,10^. Recently, attention has focused more on how the physical properties of objects and their lighting environment might give rise to a metallic appearance^11–13^. In the present study, we simulated metal objects through physically-based rendering, which varied according to two physical properties: metal smoothness, and the bumpiness of a transparent coating. These properties were chosen, not because they are interesting in and of themselves (although the visual properties of both surface and coating properties have high economic value), but because they allow us to indirectly and reliably manipulate the low-level image statistics of the stimuli, as we explain later in the paper. This allows us to interpret psychophysical data through image analysis, and the performance of corresponding computational models, in order to connect the optical properties of materials, image statistics, and visual features. We used a quantitative paradigm of multidimensional, suprathreshold perceptual judgements—conjoint measurement—to evaluate how each property affected observer judgements of metallicity, and related this to the performance of model observers using candidate image features.

### Metallicity vs glossiness

Metallicity as a visual property of materials has received little attention when compared with glossiness, and it is worth contrasting the two. Gloss, and its associated characteristics of highlights, sheen, haze, and texture distinction, is present when incident light is reflected at the air-material boundary of surfaces, giving a ‘specular’ reflectance component, before otherwise penetrating into the material and contributing to an object’s ‘diffuse’ reflectance, or albedo. Glossiness therefore affects not just the surface colour of an object, but also the spatio-chromatic statistics of its corresponding image, with images of glossy objects sharing the chromatic signature of the illuminant despite having dissimilar surface colours. Within a natural scene, illuminants will typically be weak in chroma, giving rise to glossy highlights that are simultaneously an increase in lightness and a decrease in saturation over an object’s surface.

Metallicity can be misconstrued as an extreme form of gloss, or, in the case of coloured metals such as gold, as a simple combination of colour plus gloss^14^. However, there are notable differences between the two. First, a typical metallic surface lacks a ‘diffuse’ reflectance; all incident light is reflected at the air-material boundary, due to the electro-physical properties of metals. There is, therefore, no distinction between highlight and surface colour. Second, the proportion of incident light that is specularly reflected is far higher for metals than for glossy materials. This gives rise to far brighter regions on the surface, and is more likely to result in a perceptible reflected image of the full environment, rather than the array of highlights—illuminants divorced from their environment—frequently seen on smooth, glossy dielectric objects such as a coffee mug. Third, a metallic surface specularly reflects equally at all angles of incidence, whereas dielectric surfaces obey Fresnel reflectance, and specularly reflect to a greater degree at grazing angles.

### Physically-based computer graphics stimuli

We used physically-based computer graphics rendering to generate stimuli, which permits the precise parameterization of stimulus spaces whilst staying within a physically plausible, ecologically valid domain of images. Specifically, we model objects with a silver base-layer beneath a colourful, transparent coating. This configuration may seem arbitrary, but is in fact a common material composition; with the majority of metals being colourless, foils and surfaces that would have an otherwise plain silver appearance are frequently coated with a clear, coloured layer. A common example may be found in the wrappers for assorted confectionery. By varying the smoothness of the silver (i.e. how constrained the distribution of angles is for surface microfacets), and the bumpiness of a transparent coating, we can generate images across a wide range of metallic/non-metallic appearance, with corresponding changes in image properties. As shown in Figure 1, decreasing metal smoothness (increasing roughness) effectively causes Gaussian blurring of the reflected environment, as light impinging on the surface undergoes greater scattering. Increasing the bumpiness of the coating has an effect very similar to applying a local disarray image transform, where pixels are subject to a random displacement field of a given scale and extent. The two axes of our stimulus space are therefore analogous to an ‘Eidolon factory’^15^, but every image in the stimulus set is a natural image, albeit a synthetic one.

**Figure 1.**
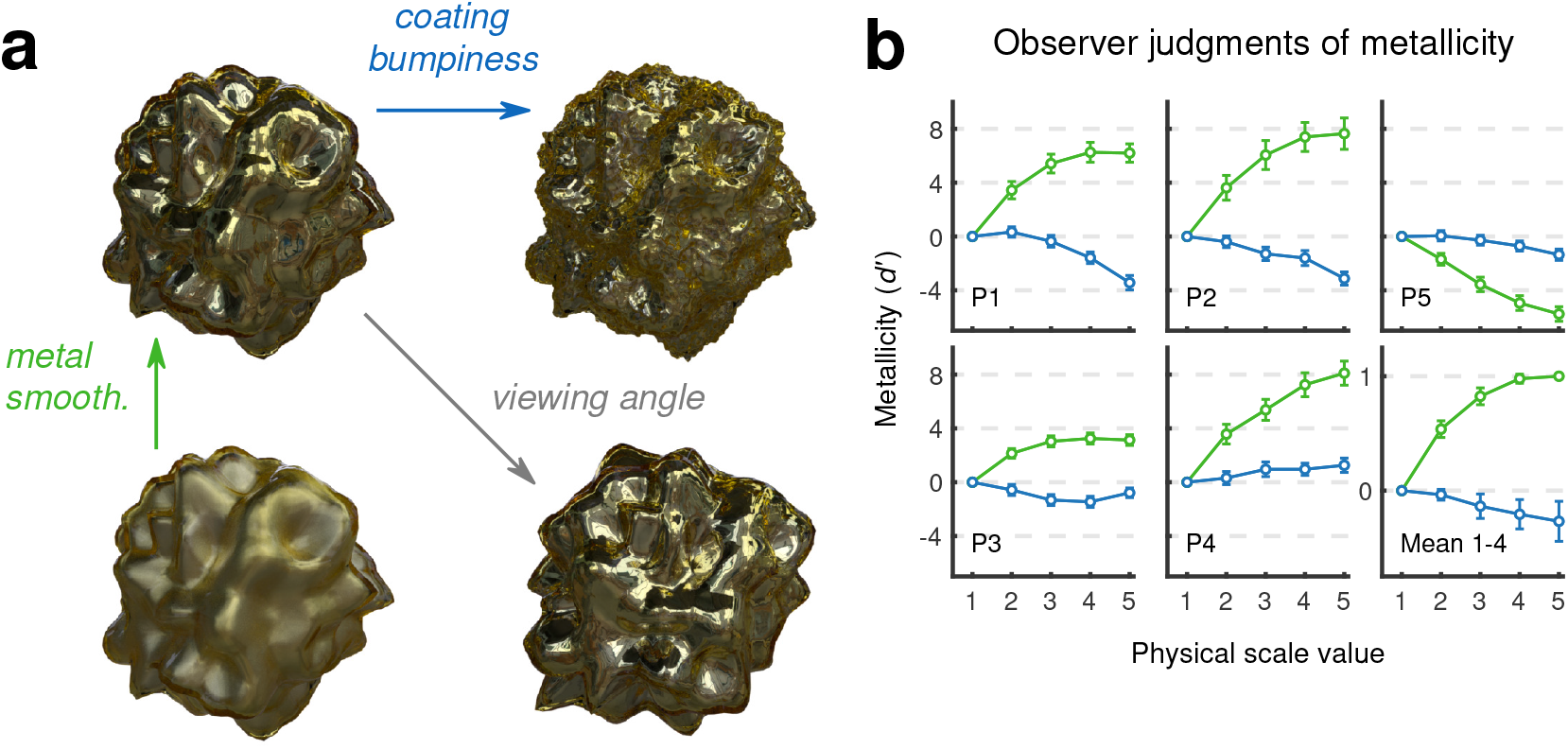
Conjoint measurement analysis of the judgment of metallicity by observers. (**a**) The parameter space used in experiments. Stimuli were physically-based rendered computer graphics of a composite coated metal object. Smoothness of the metal, and bumpiness of the coating, vary from zero to a physically plausible upper limit. The objects could be viewed from eight possible rotations around the z-axis. (**b**) Conjoint measurement plots for individual observers. Plots show the contributions of metal smoothness (green) and coating bumpiness (blue) to judgments of metallicity. Error bars for individual participants show 95% confidence intervals obtained via bootstrap. The lower right-hand plot shows the mean of normalized estimates for P1-4, with error bars showing ± s.e.m. (*N* = 4).

There are sound arguments to avoid the use of natural images in visual neuroscience, due to their complexity and unwieldiness^16^. However, manually programmed synthetic stimuli (e.g. sinusoidal gratings, plaids etc.) can only be useful insofar as the ‘recipes’ for particular percepts (or neuronal activations) are known; for material perception it is paramount that stimuli appear as convincing examples of real life materials, and using synthetic natural images, obtained with physically-based rendering^17^, allows experimenters to investigate high-level visual percepts with precision and control. There is neurophysiological evidence from both humans and nonhuman primates that material qualities and judgements may be represented in higher cortical areas, particularly in medial regions of the ventral extrastriate cortex, such as the inferior extrastriate area^18^, collateral sulcus^19,20^, and the region extending from the fusiform gyrus into the collateral sulcus^21^. Gloss-selective neurons have also been found in the inferior temporal cortex of macaque monkey^22^. As Rust and Movshon say themselves, “for neurons with complex properties whose circuitry is unknown (such as those in higher cortical areas), these methods [exploratory experiments with natural stimuli] may be the best or even the only way to begin”. For metallic objects, this necessitates their embedding within a (natural) environment, even if they are to be displayed without any background, as in this study. For this reason, we rendered all objects within a spherical light probe of a natural environment.

## Results

### Observer judgment of metallicity

To evaluate how observer judgments of metallicity varied throughout the stimulus space, we carried out a conjoint measurement task. This experimental paradigm attempts to model the trial-by-trial decisions of an observer, and so provide a quantitative measure of suprathreshold perceptual judgments as a function of stimulus properties. Each trial comprises two objects, with each object having a determined value for each of the two dimensions of the stimulus space: metal smoothness and coating bumpiness. The task required each observer to respond to 1300 trials, indicating which object within a pair was more likely to be made of metal. Rotations of objects were randomised, and never the same within a single trial, to prevent direct (i.e. pixel-based) comparisons. We consider this a minimum criterion of robust material perception—that the same object within the same environment should appear to have the same material properties, independent of viewing angle or rotation of the object within the environment. Given the pattern of responses and their corresponding trial indices (the smoothness and bumpiness levels of each object), a conjoint measurement model is fit to the data by maximum likelihood estimation (see the methods section for more details). For all observers, both dimensions of the stimulus space had a statistically significant effect on metallicity judgments (*p* < 0.001), as evaluated by performing likelihood ratio tests on the independent (only one dimension effects judgements) and additive (both dimensions have an effect but do not interact) nested hypothesis models. The conjoint measurement plots for five observers are shown in Figure 1B.

### Image statistics

Little is known about the visual features and statistics of images that drive the perception of metallicity. From a physics standpoint, metals owe their appearance to a uniform specular reflection over their entire surface, and their exceptionally low absorptivity in the visible region of the spectrum (with the exception of chromatic metals such as gold). Recent work has related metallic appearance to rendering parameters such as surface smoothness and the quality of illumination^11^. While this does inform on which properties of a scene give rise to metallic objects, it does not explain why they look the way they do to observers. To address this question, we developed computational models with which to simulate the conjoint measurement task, testing to see if low-level image statistics could provide a basis for replicating the performance of observers. Although we are primarily concerned with image correlates for metallicity, we first sought models that could discriminate variations in metal smoothness and coating bumpiness—the two physical properties we varied in the conjoint measurement analysis.

#### Spectral analysis

The relationship between image statistics and material appearance has received much attention in recent years, and the use of global statistics, such as intensity histograms, has proved controversial^23–28^. The role of spatial frequency analysis has been implicated for texture appearance^29^ and in particular has proved useful in accounting for fabric appearance^30^; here we adopt a similar approach.

Power spectra were calculated for variations in metal smoothness and coating bumpiness. Images were first windowed to minimise edge effects when computing the Fourier transforms. Spatially two-dimensional images give rise to two-dimensional spectra in frequency space; these were interpolated from *x*- and *y*-axis frequencies onto a one-dimensional radial (*r*-) frequency axis, equivalent to averaging frequency spectra by orientation as per convention^31^. As with studies into the statistics of natural images, power spectra showed a slope of approximately 1/ *f* when plotted on log-log axes^31–33^. As seen in Figure 2A, metal smoothness had a strong effect on this slope, with greater roughness attenuating higher spatial frequencies in the image (Fig. 2A, left). This was well conserved across the eight rotations of the object (Fig. 2A, right), with good agreement in the mean spectral power over the high-end of the spectrum (between 23 and 53 cycles per image, or cpi, as given in previous studies^30^) for particular levels of smoothness. Coating bumpiness also affects the power spectra of images (Figure 2.b); the divots and bumps of the coating create distortions in the images where previously image regions were fairly uniform. This results in low-frequency components being slightly attenuated, as the coating disrupts the regular curvature of the objects. However, there is a poor concordance of spectra for individual levels of bumpiness across the different rotations of the object. Surprisingly, increasing coating bumpiness results in negligible increases in the power of high-frequency components, despite the apparent increases of fine details in the image. The amplitude spectrum, then, is not a good candidate for reliable judgments of coating bumpiness.

**Figure 2.**
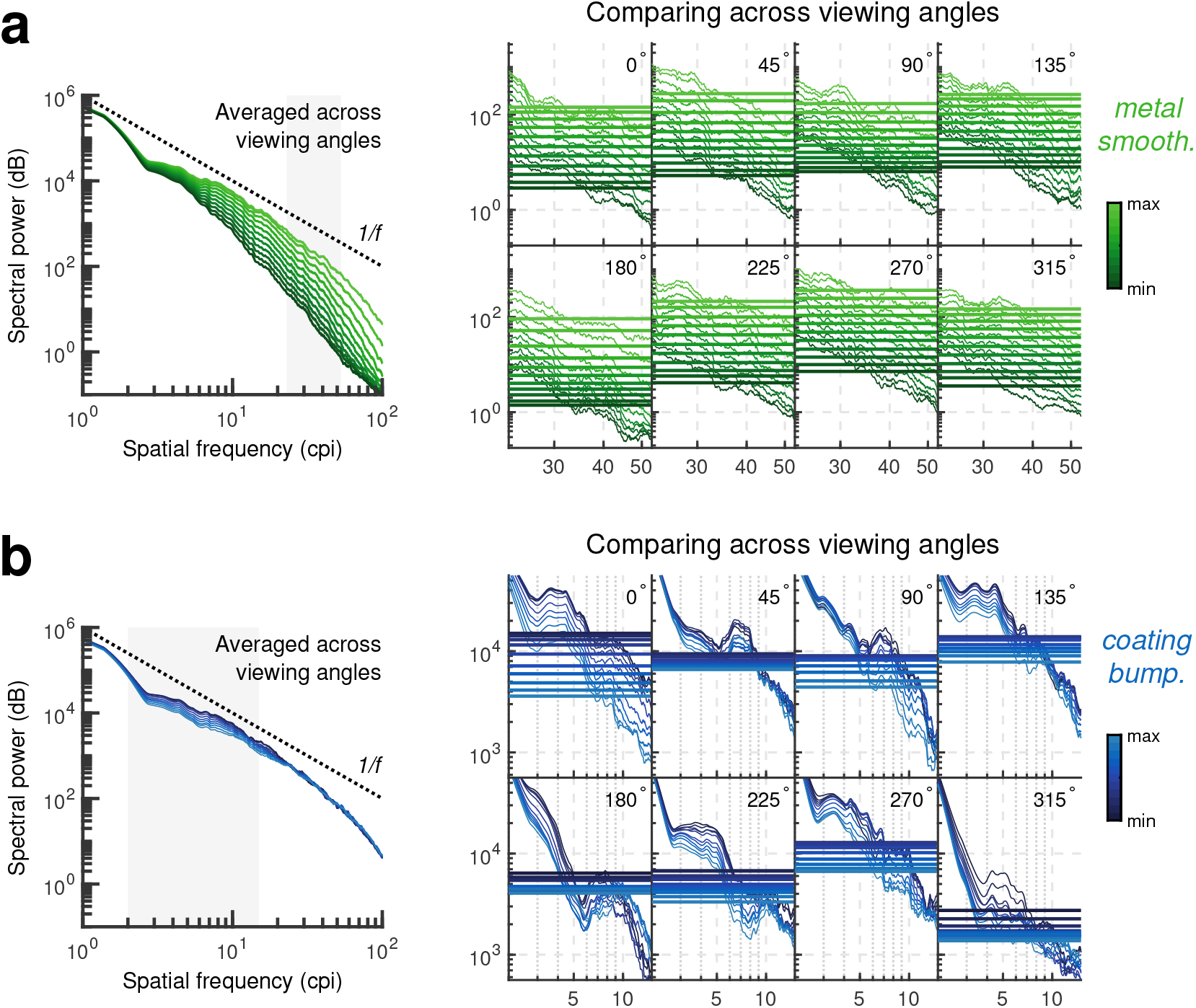
Spectral analysis of the stimuli used for conjoint measurement. Power spectra are shown for varying levels of (**a**) metal smoothness (green scale bar) and (**b**) coating bumpiness (blue scale bar) of the stimuli, interpolated onto a one-dimensional scale of radial frequency. Left: Spectra have been averaged across all eight viewing angles of the objects. Right: The power spectra for each individual viewing angle are shown separately, over the region shaded in grey in the left-hand plots. The mean power over the range shown is given by a horizontal line.

An immediate evaluation of whether spatial frequency distributions can account for material appearance can be made by swapping the Fourier amplitude spectra or phase responses of images of different materials^34,35^. It is clear from amplitude-phase swaps (not shown) that metal smoothness is primarily determined by amplitude spectra, while coating bumpiness is determined by both amplitude and phase spectra.

#### Steerable pyramid analysis

Although global spatial frequency features can sometimes provide good metrics to evaluate texture features, in particular when images are of a single object and viewpoint, and other material features are kept constant, they perform poorly outside limited cases. Another image processing approach, which is more robust against changes of these nature, is to use steerable pyramids^36–38^. Steerable pyramids are constructed by computing the outputs of oriented linear (Gabor-like) filters, at different spatial scales of an image. Algorithmically this is achieved by iteratively blurring and downsampling the image, computing filter outputs at each level, as well as marginal statistics over the images and residuals such as their variance, mean, and kurtosis. At a basic level, this method can be used to map local orientations throughout an image, obtaining an ‘orientation field’^39^, and is sensitive to both local and global image features. Because autocorrelations and correlations across scales and orientations are represented in the pyramid, the resultant catalogue of parameters captures many of the spatially invariant statistics and features of textures, most directly demonstrated by the *de novo* synthesis or the extrapolation of visibly similar textures^40,41^. While this catalogue may contain several hundred parameters, they are retained in a comprehensible, readable format, and are suitable for image analysis. While steerable pyramids are no longer the state-of-the-art in texture synthesis, having been supplanted by convolutional neural networks featuring many thousands or even millions of parameters^42^, they remain useful to vision science in their ease of implementation, interpretability (i.e. they are not a ‘black box’), and similarity in principle to the processing understood to occur in the human visual system. Recently, steerable pyramid outputs have been shown to correlate with patterns of neural activity in the macaque visual cortex in response to images of textures^43,44^.

The synthesis side of steerable pyramid analysis provides an alternative to Fourier phase-swapping. Images may be synthesised starting with either noise or a source image, and transferring the steerable pyramid parameters from a desired template pyramid. Starting with an image taken from the center of the parameter space as the base image allows us to evaluate if the steerable pyramid parameter catalogue captures the necessary image statistics to describe (and synthesise) the wider parameter space, and to probe how different components of the parameter catalogue contribute to each parameter’s associated percept. As shown in Figure 3A, which contains the central image of the stimulus space in the middle, and four synthetic images around it, steerable pyramids can sufficiently capture the properties of the stimulus space for both metal smoothness and coating bumpiness.

**Figure 3.**
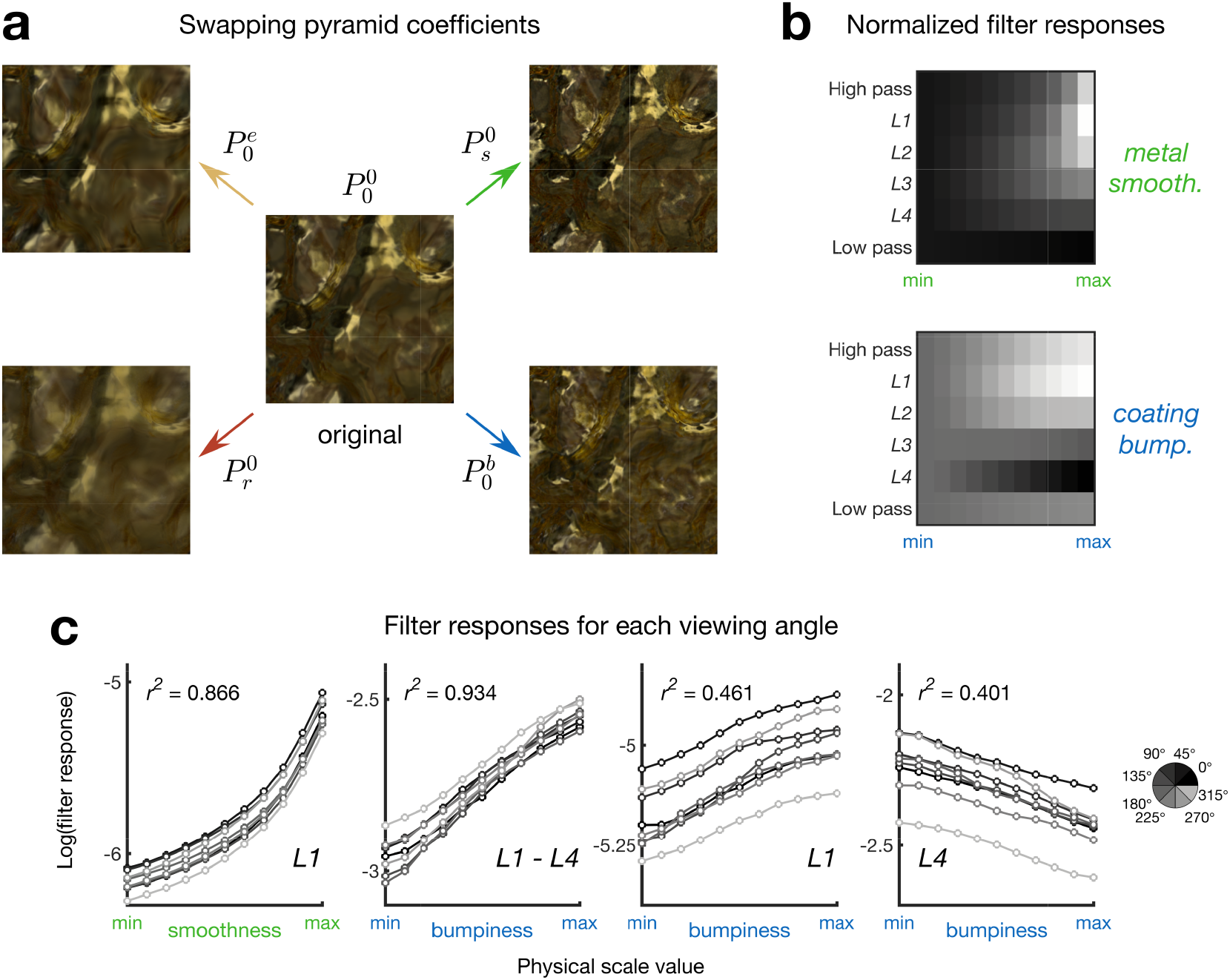
Steerable pyramid analysis of the stimuli used for conjoint measurement. (**a**) Steerable pyramid synthesis, starting from an image at the center of the stimulus space 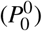. The synthesis process attempts to match the pyramid parameters of that image with the parameters of another pyramid, in this case from four different sides of the stimulus space, reaching smooth metal 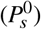, rough metal 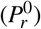, even coatings 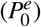, bumpy coatings 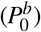. (**b**) Heatmaps showing filter responses summed over orientations, for each level of the steerable pyramid. Values of each level have been normalized to the lowest physical scale value, to show the relative changes throughout the stimulus space, averaged across all viewing angles of the objects. (**c**) The response of Level 1 filters is a good predictor for metal smoothness, while the comparison of Level 1 and 4 filters is a good predictor for coating bumpiness, across all viewing angles. For each plot, individual data series are for a single viewing angle.

Images in the stimulus set were analyzed with a steerable pyramid of four layers and four orientations. The responses of linear filters at different spatial scales, averaged over filter orientations and across all object rotations, are given as heatmaps in Figure 3B. All values presented are logarithmically transformed, as per the typical responses of neurons in the early visual pathway, although this did not effect results. As expected (and in concordance with the results of spectral analysis), increasing smoothness correlated with an increase in the magnitude of responses for filters at low levels of the pyramid, corresponding to the presence of fine details in the image. Increasing coating bumpiness also correlated with a clear increase in these responses. This corresponds with the fine edges and corners that coating bumpiness introduces in the images, which are poorly described by the global spatial-frequency analysis shown in Figure 2. Outputs of the filters for the fourth level of the pyramid reduced with increasing coating bumpiness, corresponding to the weakening of low-frequency components, which is also seen in Figure 2B. While both fine (high spatial frequency) features and broad (low spatial frequency) features are directly affected by coating bumpiness, neither one of these measures is directly comparable across different object rotations, and a visual system employing one of these metrics alone to estimate material properties would therefore exhibit limited viewpoint invariance, a requirement for robustness. However, by comparing the activations of linear filters across different scales, namely by computing the ratio of Level 1 outputs to Level 4, a metric is obtained that shows strong correlation with coating bumpiness both within and across viewing angles, shown in Figure 3C.

To see whether both axes of the conjoint measurement stimulus space could be judged reliably using the outputs of steerable pyramid analysis, we performed a difference scaling task with models using Level 1 and the difference between Level 1 and Level 4, to evaluate metal smoothness and coating bumpiness, respectively. The results, shown in Figure 4A, reflected the scaling functions of observers, which had previously been measured in a Maximum Likelihood Difference Scaling (MLDS) task to find an equidistant spacing for the conjoint measurement stimulus set (further details found below in the Methods section).

**Figure 4.**
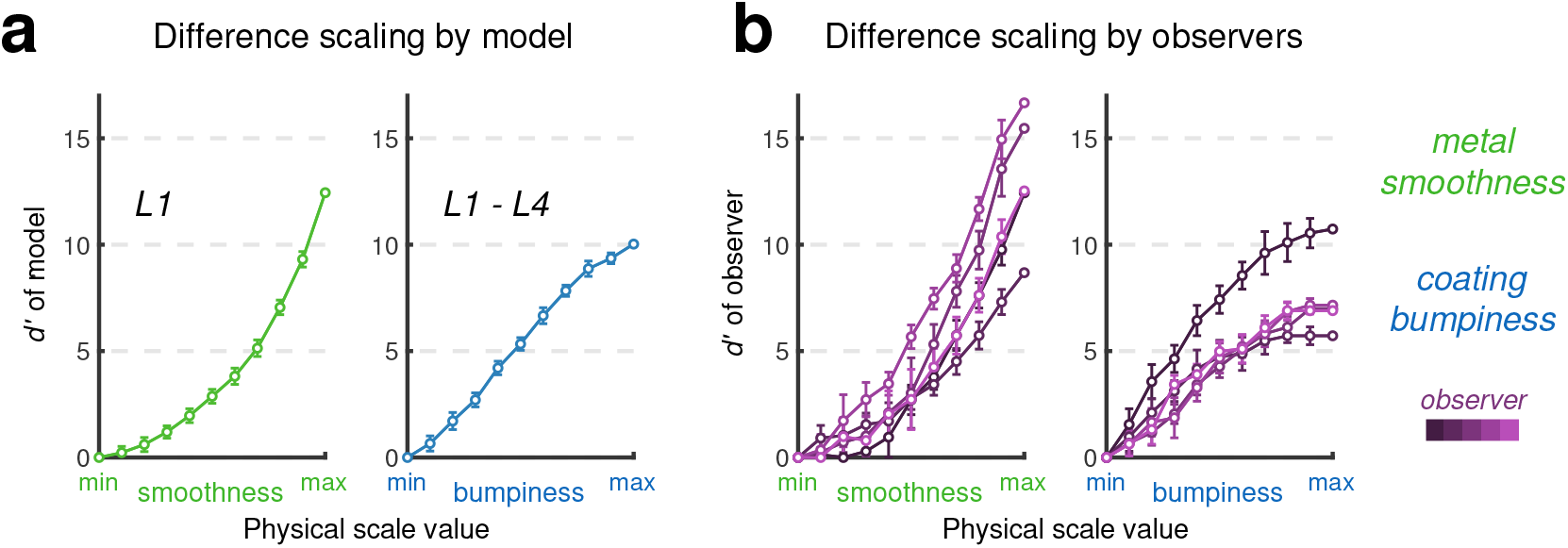
Comparison of computational models and observers for difference scaling of the dimensions of the stimulus space. (**a**) Simulated difference scaling using models of metal smoothness and coating bumpiness. The models use only the output of steerable pyramid levels for comparing trials. (**b**) Difference scaling results obtained from 5 observers completing an MLDS task. Error bars show 95% confidence intervals obtained via bootstrap.

**Figure 5.**
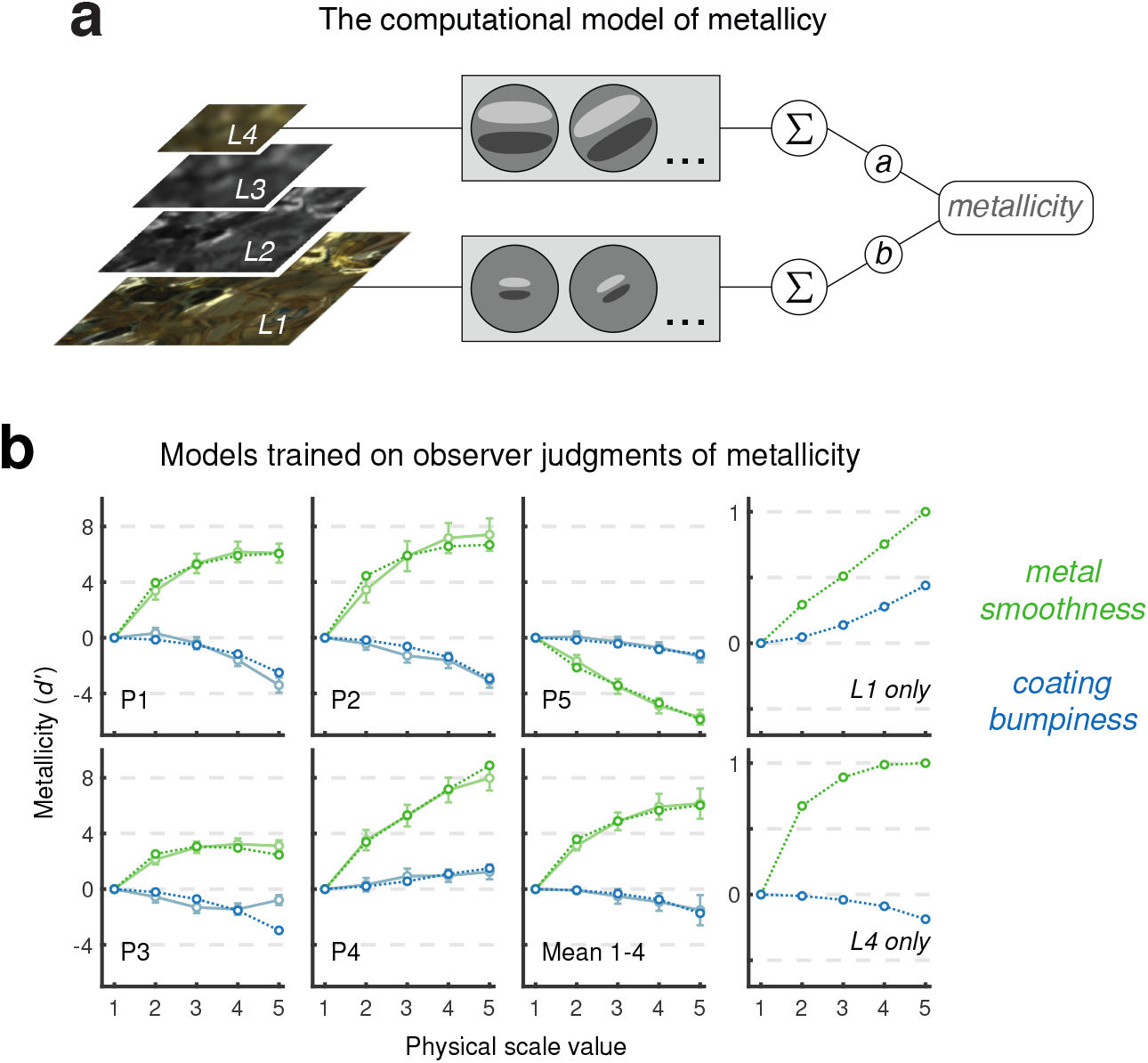
Modeling observer judgments of metallicity. (**a**) The computational model of metallicity, taking a weighted comparison of the outputs of Level 1 and Level 4 of the steerable pyramid decomposition. The pyramid is shown on the left, with higher levels resulting from applying a Gaussian blur to preceding levels and downsampling the spatial resolution by a factor of 2 (not to scale). The same linear oriented filters operate on all levels, such that lower levels generate responses owing to an image’s fine details, and higher levels generate responses owing to coarse features. The model has only two free parameters, *a* and *b*, the weightings of responses for Level 4 (coarse features) and Level 1 (fine details). (**b**) This model accounts for the observer judgments of metallicity. Conjoint measurement contributions are shown for metal smoothness (green) and coating bumpiness (blue). Observer estimates obtained via MLCM are shown in faded colors as in Figure 1, with computational model fits shown in dotted lines. Also shown are the conjoint measurement contributions for computational models using only a single level of the pyramid. Level 1, corresponding to fine image features, at the top right, and Level 4, corresponding to coarse image features, at the bottom right. Error bars for pooled participant data show ± s.e.m. (*N* = 4) and for individual observers show 95% confidence intervals obtained via bootstrap.

### Modeling metallicity

We then sought to model the performance of observers in the conjoint measurement task, where they made judgments about the relative metallicity of pairs of stimuli that varied in both metal smoothness and coating bumpiness. In keeping with the literature on gloss, which suggests that glossiness in natural images may be inferred by (or at least correlated with) global luminance distributions, we compared a range of models that used different estimators of global image statistics as a predictor for metallicity, summarised in Table 1. Skewness (abbreviated to Sk in Table 1) of the luminance distribution (histogram) has previously been found to correlate with the perceived glossiness of materials in images^23,24^. A model using luminance distribution skew showed an increase in metallicity judgment with increasing metal smoothness of the metal base, as seen in the observer data. However, this model was not affected by coating bumpiness, whereas all of the observer data sets were best fit by additive models rather than independent (i.e both dimensions of the stimulus space have an effect on observer judgments). Models estimating metallicity according to global contrast, such as dynamic range (either simply taking the ratio of the brightest and dimmest pixel, DR, or a more robust average of the brightest and dimmest 10 pixels, rDR) or Michelson contrast (MC) gave similar results, with predicted metallicity varying only as a function of metal smoothness.

**Table 1.**
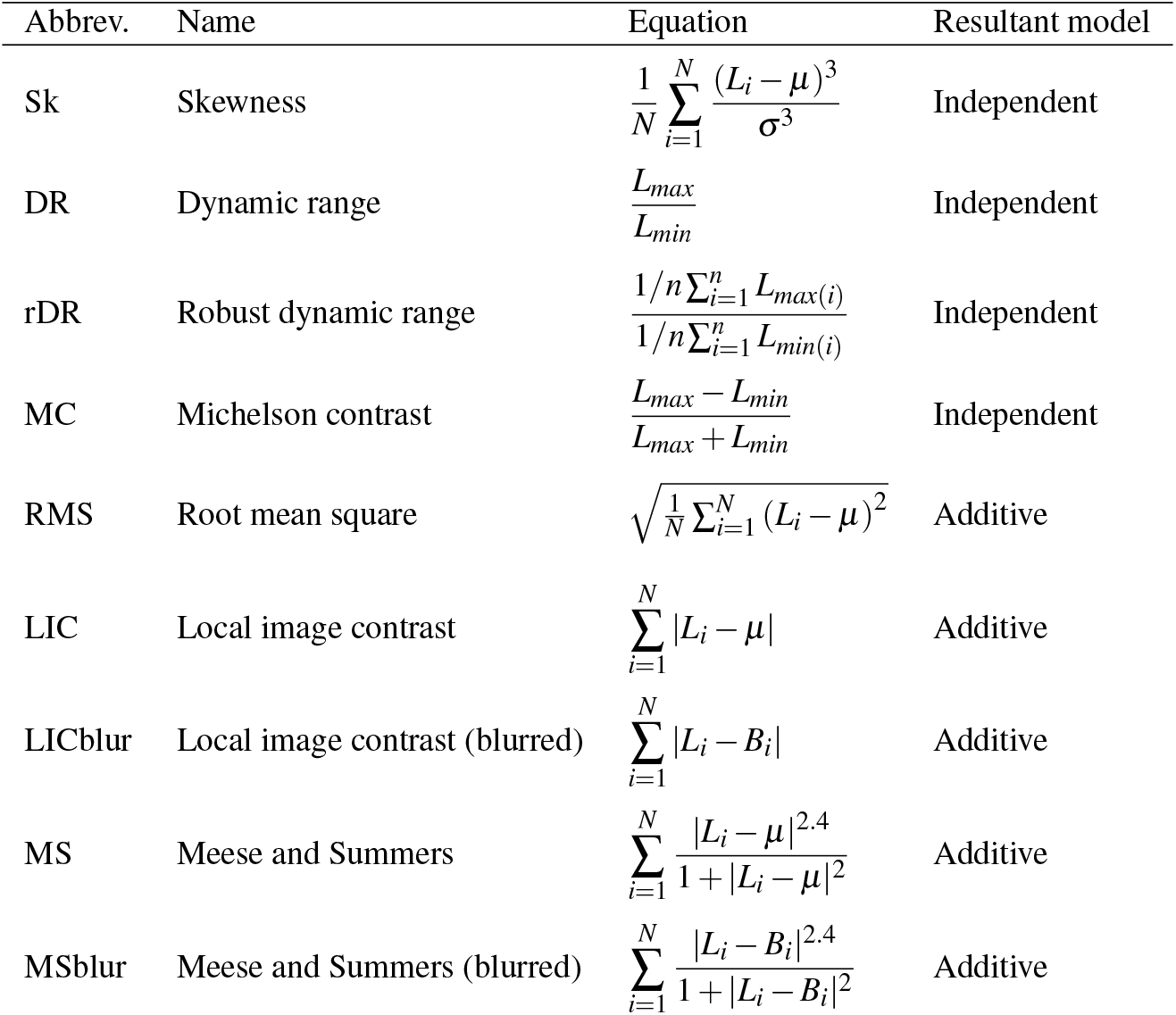
Table summarising the global contrast estimators used in modelling conjoint measurement assessment of metallicity. *N* denotes the total sample number (i.e. the number of pixels in the image, while *n* denotes the number of a sub-population. *B* is a highly blurred copy of the original luminance image, *L*. *μ* is the mean luminance value and *σ* is the standard deviation of the luminance distribution.

A model using root mean square (RMS) as a global contrast measure gives rise to results highly concordant with the observer P1-4 average. RMS has an established record as a reliable indicator of image contrast^45^, however, as Bex and Makous remark, a global metric such as RMS is difficult to relate to models of the human visual system. A method of estimating local image contrast (LIC), by simply averaging differences between each pixel and the image mean (for the region containing the object) gave similar results, with metallicity increasing monotonically with metal smoothness, and slightly decreasing with coating bumpiness. If differences were computed not against the image mean, but rather a local mean (the same location in a heavily blurred image, LICblur), the model also gave similar results to participant data. We also implemented the Meese and Summers method for estimating global contrast with gain control^46^, which has recently been found to more accurately predict observer performance in contrast estimation tasks^47^. Models employing this estimator, using either the mean of the image (MS) or pixels in a blurred image (MSblur) to calculate local contrast, performed similarly to the observer P1-4 average.

While some of the models we tested gave rise to estimates of metallicity similar to those obtained from observers in the conjoint measurement task, we also considered a model that computes global contrast directly from the outputs of steerable pyramids, shown diagrammatically in Figure 4A. This is in keeping with an established, earlier ‘generalized multiple-channel model of visual detection and perception’^29,48,49^. We wanted to see whether this simple computational model, taking as inputs the linear filter responses across different spatial scales, could replicate the performance of observers during the conjoint measurement of metallicity. In particular, it was of interest whether such a model could accommodate the individual differences we found in observers, as well as the average of similar observers (P1-4). A model was built that took weighted responses of just two different levels of the steerable pyramid—Level 1 (corresponding to fine details in the image) and Level 4 (corresponding to coarse image features). As shown in Figure 4A, the model has only two free parameters—the weightings (taking any real value) for each level. As Figure 4B shows, this two-parameter model can give very similar results to observer data, when the weightings of each level are optimised for each observer’s conjoint measurement model. The same figure also shows how a model performs that is using only the outputs of Level 1, that is, a model that is essentially estimating smoothness differences as a proxy for metallicity. This model performs similarly to P4 and the inverse of P5 (ie. P4 could be equating smoothness with metallicity, while P5 could be equating roughness with metallicity), while the conjoint measurement models of other observers agree more with a primarily Level 4-based model. This is also apparent by examining the optimised weights for the computational models fit to each observer, shown in table 2, as for both P4 and P5 the optimal weighting for Level 1 has a greater magnitude than Level 4. These individual differences will be addressed in the following section.

**Table 2.** The optimal weightings of Level 1 and Level 4, for a computational model of metallicity fit to observer data.

| Observer | L1 | L4 |
| --- | --- | --- |
| P1 | 0.262 | 13.955 |
| P2 | 0.110 | 15.939 |
| P3 | -1.832 | 11.514 |
| P4 | 5.527 | 4.988 |
| P5 | -3.852 | -2.691 |
| Mean 1-4 | 1.017 | 11.600 |

## Discussion

Investigating material perception presents some particular challenges to visual neuroscientists and psychophysicists. Stimuli require both verisimilitude and versatility; they must be convincing representations of the material percept in question, and they should have properties that can be predictably and independently varied. Our stimuli, of physically-based rendered computer graphics, meet these requirements. The model of a coated metal object, with varying metal smoothness and coating bumpiness, allows for a continuous stimulus space extending across both metallic and plastic appearances, and every location within this space is a (synthetic) natural image. Additionally, we have shown that these physical properties predictably determine image statistics of the stimuli, through a steerable pyramid analysis, which are then directly relatable to the percept of metallicity.

Most observers tended to judge smoother metal as more metallic, confirming a previous finding for ambient lighting conditions^11^. However, one participant (P5) showed the opposite effect, raising questions over the homogeneity of the population of observers for metallicity judgements, and material perception in general. This double potential for metallic appearance, whilst at first seeming contradictory, is consistent with the physical properties of metals. Specifically, the hardness of metals enables their surfaces to stably retain both a highly polished, mirror appearance, when microfacets are approximately coplanar, as well as a roughened, matte appearance, when microfacets are highly misaligned. Indeed, it is reasonable to assume that visual material judgements are heavily influenced by each observer’s typical environment, in this case whether they are more likely to see matte metallic surfaces such as anodized aluminium electronics, weathered coins, or brushed metal appliances etc., rather than highly-polished and mirror-finish objects. Individual differences are neatly captured by our computational model of metallicity by re-weighting the levels of the pyramid used in the model, as summarised in Table 2.

As increasing metal roughness tends to decrease the strength of metallic appearance for the majority of observers, this may initially suggest that observers are simply estimating the degree of articulation within the reflected image of the object’s environment, and relating that to metallicity. Rougher metals blur specular reflections, attenuating high spatial-frequency components as seen in Fourier power spectra. However, the negative contributions to metallicity of coating bumpiness, which actually increases the degree of articulation over the object’s surface (as evidenced by increasing the activations of lower levels of steerable pyramids), suggests a different mechanism. Two possibilities remain when the experimental results are considered in the light of the modelling we have undertaken. It could be that fine (high spatial-frequency) details and edges are less important to observers than broad (low spatial-frequency) details. Coating bumpiness introduces additional fine details, such as edges and corners, while disrupting gradual, coarser features, and this is particularly the case when the reflected environment image is already blurred (in the rougher regions of the stimulus space). Or, it could be that observers are selectively relating fine details to metallicity only when they are recognised as part of the reflected image of the environment, whose edges are generally continuous and extended owing to the properties of natural images. While coating bumpiness increases the fine details on the object’s surface, these show a reduction in collinearity and co-circularity (i.e. correlations between neighbouring oriented filters of the same orientations). It is plausible that observers are sensitive to this (e.g.^50^), and that the resultant irregularly wrinkled appearance does not contribute to perceptions of metallicity in many observers, and perhaps may even disrupt it.

If observers have built more of a matte metallic association, ‘shiny’ objects’ appearances may be more readily associated to plastic-coated objects, or—and particularly if seen without a background for context—to transparent glass objects. As objects in our stimulus set were presented in a void, one initial question the perceptual system may have to answer is whether they are reflecting light or transmitting light—are the objects mirrors or windows? For a perfectly polished metal surface, which reflects a mirror image of a given environment, there exists a corresponding transparent object of another environment, which gives rise to a very similar image for the observer. Tamura et al. recently investigated this ambiguity in material perception, and found that without motion cues from dynamic stimulus presentations, observers classify mirror objects as glass between 8 and 40% of the time, depending on the object geometry^4^. This study only considered perfectly smooth surfaces, but we can speculate as to how increasing roughness might improve or disrupt classifications. When light reflects off a rough metal surface, microscale geometric inhomogeneities have the effect of scattering a fraction of the light at angles other than the angle of reflection, the result being a blurring of the image. For rough transparent objects, light is scattered at both entry and exit points. Rather than simply blurring the transmitted image, this also tends to flatten and desaturate it, giving a ‘frosted’ or translucent appearance, as light from illuminants may be distributed to any region of the object after the initial scrambling at entry. This effect is shown in Figure 6, where silver and glass objects are rendered with different levels of physical roughness. For silver and glass objects whose roughness gives a similar blurring of the reflected environment, glass objects show a more tightly constrained luminance distribution, lacking the extreme highlights and shadows present over the surface of the metal object. We would therefore expect that observers who confuse smooth metal for glass might be less prone to this error for objects whose surfaces are not perfectly smooth.

**Figure 6.**
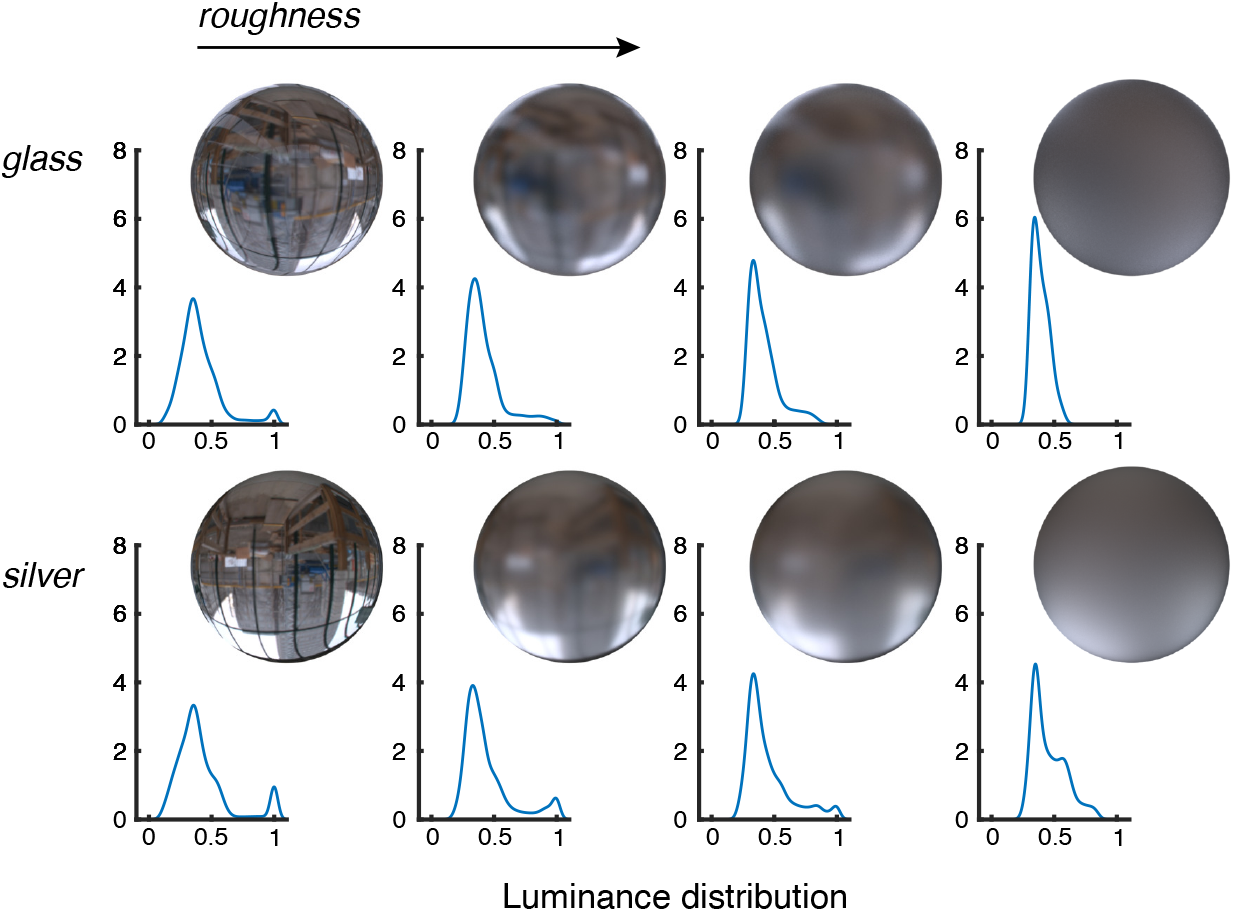
Comparing the effect of roughness on glass and silver objects. Glass (top row) and silver (bottom row) spheres have been rendered with varying physical roughness, increasing to the right. The objects have a similar luminance profile when completely smooth, making it hard to discern which material is which. However, increasing roughness has a more pronounced effect on the glass object than the silver object, as the luminance distribution is more tightly constrained, despite a similar apparent level of image blurring.

In this study, stimuli have consisted of objects devoid of any background, although it is almost certainly the case that in everyday vision, observers are heavily influenced by environmental context in their appraisal of metallic surfaces. In such cases, the relative luminance of surfaces with respect to environmental illumination is likely one of the most reliable potential cues, as typical metallic surfaces reflect all of the incident light falling on them. This may be particularly critical for the perception of very rough, even matte, metal surfaces such as anodized aluminium, or patinated metals. The surfaces of such materials cannot be said to be devoid of a ‘diffuse’ appearance, but rather fall along a continuum of matte and mirror appearance. It is therefore also possible that there is not one single visual feature that confers the appearance of metallicity, but rather several categories e.g. polished, matte, painted, rusted etc.

## Conclusion

Using physically-based rendering to generate synthetic natural images, we investigated the relationship between observer metallicity judgements and low-level image features. While global image features such as frequency spectra, dynamic range, and luminance contrast were unable to account for robust observer judgements of the metallicity of stimuli, local image features—the activations of Gabor-like filters computed with steerable pyramid analysis—could be used to replicate observer performance, including significant individual differences.

## Methods

### Stimuli

#### Scene modelling

Stimuli consisted of computer-generated images, the primitive of which was a metal sphere randomly deformed to contain smoothly curving projections, enveloped within a coating. This shape exhibits several required properties. It has a wide array of curvatures, including regions of both highly positive and negative curvature, as well as numerous saddles, which ought to aid in visual judgements of material qualities^51^. This being so, the shape has a distinct solidity, with flatter regions of the structure preventing it being perceived as a smoothly flowing or rippling liquid. In combination, the flatter and more curvaceous regions of the shape give rise to specular reflections of the environment across the gamut of those seen day-to-day in metallic surfaces, such as cars, cutlery, and cookware. While it has rotationally invariant statistical features—the height and frequency of projections—these are distributed randomly, ensuring that when viewed from different angles observers must make a material judgment, rather than a direct image comparison. Direct image comparisons (i.e. comparing the same region of the environment reflected in two or more objects) are also hindered by the high frequency of random projections, which scramble the locations of the environment’s features.

Stimuli were modelled using Blender 2.77, open-source 3D computer graphics software, in a similar manner to a recent study on metal and glass visual perception^4^. First, the object was created as an icosphere with two subdivisions. Then, the mesh is subdivided, with the number of ‘cuts’ set to two, ‘fractal’—a measure of random deviation in the mesh—set to 10, and ‘along normal’ set to one, ensuring projections lead directly out from the center. The ‘subdivision surface’ modifier is then applied with two subdivisions, and smooth shading was specified. The coating is defined as the same shape scaled up by a factor of 1.05. This has a further two subdivisions applied, giving a resolution that permits smooth bumpiness. The bumpiness is defined with a ‘displace’ modifier, configured with a Stucci texture of size 0.05 and turbulence 1.00. The ‘strength’ level of the modifier varies between 0 and 0.05. As the name may suggest, the ‘Stucci’ procedural texture provides a random displacement field that evokes a decorative process, as opposed to naturally occurring or unrealistic options such as Voronoi patterns or simple spheres.

#### Rendering

Physically-based rendering was carried out with Mitsuba^52^, configured hyperspectrally with 31 10-nm wavelength bins between 395 and 705 nm. Rendering specifications (materials, rendering engine, integrator etc.) were defined with RenderToolbox where possible^53^. The object was assigned a ‘rough conductor’ BRDF with the spectral reflectance of silver. Roughness (the inverse of smoothness) was varied between 0 and 0.15, using the ‘ggx’ microfacet distribution model, as it has been found to give more physically accurate results over the standard Beckmann distribution^54^. The coating was assigned a ‘dielectric’ BRDF with a refractive index of 1.4, a plausible value for a varnish or glaze, and an interior ‘homogeneous medium’ with absorbance defined as 0, 0.05, 0.60 in RGB (which is then interpolated to hyperspectral data by Mitsuba), at a ‘scale’ of 20. Eight renders of each coated object were obtained by rotating at 45° intervals around its vertical axis.

Objects were illuminated by the ‘Overcast day/building site’ environment light probe made by Bernhard Vogel (http://dativ.at/lightprobes/), which provides a realistic environment that does not require tone mapping or compression of the luminance space for presentation on a standard display. Path tracing was computed with the ‘extended volumetric path tracer’ with an infinite maximum path length. The ‘hide emitters’ function created images of objects in a black void. The ‘low discrepancy sampler’ was used; a sample count of 256 was found to give images of sufficiently low noise. For the MLDS stimuli sets, 176 images were rendered over 17 hours on a Macbook Pro (early 2015). For the MLCM stimulus set, a further 200 images were rendered over over 20 hours.

Rendering outputs were 31-dimensional 512 × 512 pixel hyperspectral images. These were transformed to LMS values based on Stockman and Sharpe 2° cone fundamentals before conversion to RGB for display on a calibrated cathode ray tube (CRT) display. The final images were 512 × 512 single precision floating point matrices.

### Psychophysics

#### Participants

Five practised observers were recruited from the Department of Experimental Psychology at the University of Oxford. All participants had normal colour vision (as determined using an HRR test) and normal or corrected-to-normal visual acuity. Participant age ranged from 23 to 28 years. All participants gave informed consent. The protocols of the study were approved by the Medical Sciences Interdivisional Research Ethics Committee at the University of Oxford, in accordance with the Declaration of Helsinki.

#### Difference scaling

In order to determine the spacing of stimuli, we initially performed a maximum likelihood difference scaling (MLDS) analysis for each dimension of the stimulus space^55^. This indirect method has advantages over more traditional appearance-based (suprathreshold) methods such as Thurstonian scales and just-noticeable difference estimation, and may arrive at more accurate estimates^56^. MLDS has been used to evaluate colour perception^57,58^, depth perception^59^, image quality^60,61^, emotion^62^ and gloss perception^63^. By obtaining approximately equivalent spacings throughout the stimulus space, we safeguard against nonlinearities and prevent one dimension from dominating the other during the conjoint measurement task.

#### MLDS procedure

Participants viewed all stimuli in a dark room on a CRT monitor (NEC, FP2141SB, 21 inches, 1600 × 1200 pixels) controlled with ViSaGe MkII (Cambridge Research Systems), which allows 14-bit intensity resolution for each phosphor. Gamma correction was performed with a ColorCAL MkII colourimeter (Cambridge Research Systems) and spectral calibration was performed with a SpectroCAL MkII spectroradiometer (Cambridge Research Systems). Viewing distance was maintained with a chin rest positioned 92 cm from the CRT monitor. Participants viewed the screen binocularly.

On each trial, participants were presented with a quadruple (i.e. two pairs of stimuli simultaneously presented) and asked to indicate for which pair the material composition of the objects appeared to have a greater within-pair difference. Trials comprising variations in metal smoothness, and variations in coating bumpiness, were interleaved. In either case, the values of the varying properties of the four objects on each trial were drawn from eleven possible values, such that all four objects had different values. Additionally, each image in the quadruple was viewed at a rotational angle drawn without replacement from the eight 45° intervals, preventing direct image comparison. Participants viewed all 310 non-overlapping quadruples, for variations in both metal smoothness and surface bumpiness, in a randomised order. Participants were given 3 s to respond to each trial and entered their response by pressing either up or down on a response box, selecting either the upper pair or lower pair in the quadruple.

#### Conjoint measurement analysis

Methods of conjoint measurement analysis, and in particular the fitting of models through maximum likelihood estimation, have been developed for some time in psychophysics^64,65^. Maximum likelihood conjoint measurement (MLCM) has been applied to colour^66^, the watercolour effect^67^, as well as material judgments^65,68,69^. The basis of this approach is to find the relative contributions of two (or more) independent physical dimensions to a single perceptual judgment. In our case, we are interested in the effect of metal smoothness, *S*, and coating bumpiness, *B*, on the percept of metallicity, *M*. When an observer is given the choice of two images, one of metal smoothness and coating bumpiness levels (*i, j*), the other of (*k, l*), and must decide which she finds more metallic, we can model the decision variable, Δ as:

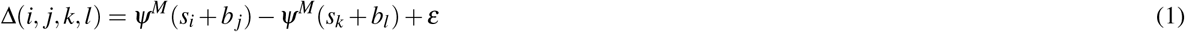

where *ψ*^*M*^ is a function that relates the physical levels of a stimulus— for example of *i*th-level metal smoothness, *s*_*i*_, and the *j*th-level of coating bumpiness, *b*_*j*_—the to the percept of metallicity. The decision variable is perturbed by Gaussian noise, 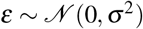.

These latent functions can be modelled with a set of nested hypotheses, with each hypothesis formulated as a generalized linear model. The most restricted model is the *independent model*, where only one dimension influences the perceptual judgment, and the observer’s decision process is uncontaminated by other dimensions. For example, if the percept of metallicity were only a function of metal smoothness, the decision variable could then be expressed as:

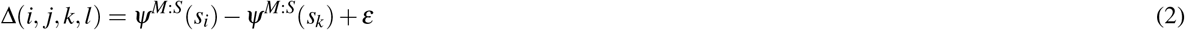

where *ψ*^*M:S*^(*s*_*i*_) is the additive contribution of metal smoothness to metallicity, computed for the *i*th-level of that dimension.

If such contamination is present, the next level of the model with increasing complexity supposes a linear mixing of the two physical dimensions, and is said to be *additive*.

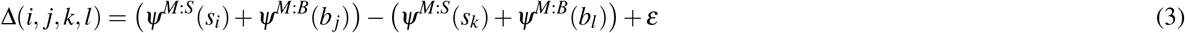

If this mixing is not linear, but the contribution of one dimension to the perceptual judgment depends on the level of the other, a fully comprehensive—or *saturated*—model is required to best account for the variance of the data. In this case, the functions relating physical levels of metal smoothness to the percept of metallicity depend on how bumpy the coating is, 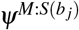, and vice-a-versa. The decision variable is then given by:

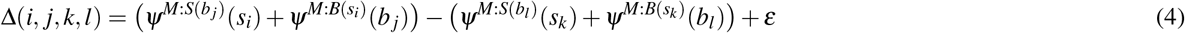

With a set of collected responses from observers, these models can be fit via optimization, to maximize the likelihood function given trial indices. This accounts for the stochasticity of observers, and allows us to compare across levels of the nested hypothesis by evaluating their goodness of fit using the log likelihood. If a more complex model results in a significantly better fit (as found by a chi-squared approximation), it is preferred over the simpler model. Ultimately we obtain values of *d*′, the sensitivity index^70^, which gives a measure of how discriminable the effect of changing physical dimensions of the stimuli is on the percept of metallicity.

#### MLCM procedure

The experiment was carried out in a similar manner as for MLDS. Each trial consisted of simultaneous presentation of a pair of stimuli, with participants asked to indicate which object appeared most likely to be made of metal. In each trial, the values of the varying dimension of the two objects were drawn at random, without replacement, from the five possible values. Additionally, each image in the pair was viewed at a rotational angle drawn without replacement from the eight 45° intervals, hindering direct image comparison. Each participant viewed all 325 pairs, in a randomised order, four times, totalling 1300 trials. Participants were given 2 s to respond to each trial and entered their response by pressing either left or right on a response box, selecting either left or right in the pair.

## Data availability

The datasets generated and analysed during the current study, along with the code required to generate the figures in this paper, are available in the Figshare repository, https://doi.org/10.6084/m9.figshare.14079807.v2. The packages for fitting difference scaling and conjoint measurement models in Matlab are also available separately at the following Github repository: https://github.com/hirschland/SupraThresh.

## Acknowledgments

This work was supported by the AHRC under Grant No. AH/N001222/1. The authors are grateful to Rafał Mantiuk for comments on an earlier draft.

## Author contributions

Both authors conceived the study. J.S.H. programmed and conducted the experiments, analyzed the data, and carried out image analysis and modeling. J.S.H. wrote the original manuscript and visualized the results. Both authors edited and revised the manuscript and approved the final version.

## Additional information

The authors declare no competing interests.

